# Lactation curve model with explicit representation of perturbations as a phenotyping tool for dairy livestock precision farming

**DOI:** 10.1101/661249

**Authors:** Ben Abdelkrim Ahmed, Puillet Laurence, Gomes Pierre, Martin Olivier

## Abstract

**Background:** Understanding the effects of environment on livestock provides valuable information on how farm animals express their production potential, and on their welfare. Ruminants often face perturbations that affect their performance. Evaluating the effect of these perturbations on animal performance could provide metrics to quantify how animals cope with their environment and therefore, better manage them. In dairy systems, milk production records can be used to evaluate perturbations because (1) they are easily accessible, (2) the overall dynamics throughout the lactation process have been widely described, and (3) perturbations often occur and cause milk loss. In this study, a lactation curve model with explicit representation of perturbations was developed.

**Methods:** The perturbed lactation model is made of two components. The first one describes a theoretical unperturbed lactation curve (unperturbed lactation model), and the second describes deviations from the unperturbed lactation model. The model was fitted on 319 complete lactation data from 181 individual dairy goats allowing for the characterization of individual perturbations in terms of their starting date, intensity, and shape.

**Results:** The fitting procedure detected a total of 2,354 perturbations with an average of 7.40 perturbations per lactation. Loss of production due to perturbations varied between 2% and 19%. Results show that the number of perturbations is not the major factor explaining the loss in milk yield over the lactation, suggesting that there are different types of animal response to challenging factors.

**Conclusions:** By incorporating explicit representation of perturbations, the model allowed the characterization of potential milk production, deviations induced by perturbations (loss of milk), and thereby comparison between animals. These indicators are likely to be useful to move from raw data to decision support tools in dairy production.

## INTRODUCTION

In the context of precision livestock farming, simple interpretive tools are required to convert raw time series datasets, now routinely recorded in animals, into useful information for on-farm decision-making. Such tools are not only expected to provide farmers with good information on performance level of individual animals, but also to detect pathological, nutritional or environmental problems affecting production traits at individual or herd scales. In dairy systems, it is well known that milk yield can be affected by events such as udder health problems [1], lameness [2], meteorological changes [3], or feed quality [4]. Such problems induce perturbations in the course of the lactation process and result in a serrated shape pattern of the lactation curve. These perturbations can be seen as deviations of the lactation curve from its typical profile. This typical profile reflects that lactation is a physiological process common to mammalian females, and as a result, its expression through time follows a general pattern [5]. It can be described in 3 phases. The first phase starts after parturition with the initial milk yield increasing to a maximum or peak yield. The second phase is a plateau-like period in which maximum milk yield is maintained for a more or less long time. The third phase is the decrease from the peak yield. This last phase can be divided into two parts according to the speed of decrease, the first one corresponding to an approximately constant declining rate of milk production after the peak yield, and the second corresponding to an acceleration of the milk yield decline as pregnancy progresses before the start of the dry period when lactation stops [6–8]. Modelling the lactation curve is a long standing issue [9] and numerous authors have proposed mathematical models allowing the characterization of milk yield dynamics*, i.e.*, the transformation of a series of temporal data into a vector of estimated parameters via a fitting procedure. The most famous and used model is the one published by Wood in 1967 [10]. The overall objective of lactation models is to reduce the variability in data by creating a profile, thereby being able to characterize an average animal milk production, or to compare the production of different animals. This strategy of using lactation models as phenotyping tool has been very useful in the past years (for instance, test-day models for genetic selection) and in a context of scarce raw data. An important limitation of these modelling approaches is that short-term perturbations are removed during the fitting procedure in order to extract an unperturbed phenotype, corresponding to a typical lactation curve. However, characterizing perturbations can be highly relevant for better understanding the resilience of dairy females regarding their milk production and therefore for making management decisions [11]. Furthermore, evaluating the effect of perturbations on animal performance could provide metrics to quantify how animals cope with their environment, and develop management strategies to find a good balance between animal welfare and performance.

The need for incorporating perturbations into lactation curve models is also driven by the development of precision livestock farming. Now, we have more frequent and reliable data and we can transition data analysis from reducing variability around average profiles to extracting variability to provide information. High throughput data has led to the development and use of statistical methods, such as smoothing methods, to capture and understand perturbations [12]. Codrea et al. [12] studied the effect of nutritional challenges on the lactation curve in dairy cows using differential smoothing procedures for quantifying biological perturbations in an animal performance. Results of this study highlighted the decline in milk yield during the challenge period for each cow, and showed the presence of other deviations with unknown causes or unrelated to the feed restriction during experiment. On the other hand, Friggens et al. [4] used a clustering procedure linked to a piecewise mixed model to characterize different responses between lactation stages and types of response for the nutritional challenges. Another study have highlighted the large differences in milk production in goats that are subject to the same diet and environmental conditions [13]. There are few other approaches to describe the shape of the lactation curves from animals faced with health problems. Lescourret and Coulon [14] had shown the huge variability of milk production in response to mastitis in both form of the lactation curve and intensity of milk production. Adriaens et al. [1] developed a novel methodology to predict quarter milk yield during clinical mastitis.

The main shortcoming of approaches cited above is the lack of an explicit representation of perturbations which are only captured through statistical objects. To overcome this limit, models on different animal species have been developed with a more explicit representation of perturbations. In the work of Revilla et al. [15] on growing piglets, a classical Gompertz equation, used to capture the unperturbed growth curve, is combined to an equation of the perturbation, used to capture the perturbation in body weight change induced by the weaning stress. Another model based on differential equations was developed to characterize the feed intake response of growing pigs to perturbations [16]. Sadoul et al. [17] used a model based on a spring and a damper to capture perturbations in physiological responses to challenges on rainbow trout. This formalism allows the characterization of perturbations with stiffness and resistance to the change of the system.

These recent modelling developments exhibit two major limits for application to lactation curve: first, they do not allow to capture multiple perturbations that may be imbricated and second they imply that the time of perturbation is *a priori* known.

In this study, we developed a Perturbed Lactation Model (PLM) that incorporates an explicit representation of perturbations and that converts individual raw time-series data into biological meaningful parameters. The fitting procedure of PLM allows the detection and the characterization of perturbations in milk time-series. The objective of the present paper is (1) to introduce the PLM model and the explicit representation of perturbations, (2) to describe the use of PLM to detect and characterize perturbations in milk yield time series with an example in dairy goats, and (3) to illustrate the role of PLM as a phenotyping tool by analyzing the variability in perturbed lactation curves on the basis of the fitting results obtained on the dairy goat dataset.

## MATERIALS AND METHODS

The PLM is composed of a lactation model, denoted *Y*^∗^, describing the theoretical unperturbed dynamics of milk yield along the lactation, and a perturbation model, denoted *π*, describing deviations from the lactation model. The list of model parameters is provided in Table 1.

**Table 1:** Model parameters

| Symbol | Definition |
| --- | --- |
| *Wood Model* |  |
| $a$ | Parameter scaling the general level of the lactation curve |
| $b$ | Parameter controlling the type and magnitude of the curvature of the lactation curve |
| $c$ | Parameter regulating the rate of decrease in milk yield after the lactation peak |
| *Perturbed Lactation Model with $N$ perturbations ($PLM_N$)* |  |
| $N$ | Number of perturbations |
| $i$ | Perturbation number ($i \in [0; N]$) |
| $t_{P,i}$ | Time of start of the $i^{th}$ perturbation |
| $k_{0,i}$ | Parameter of intensity of the $i^{th}$ perturbation |
| $k_{1,i}$ | Parameter of collapse speed of the $i^{th}$ perturbation |
| $k_{2,i}$ | Parameter of recovery speed of the $i^{th}$ perturbation |

The dynamics of daily milk yield (*Y*(*t*), in kg) during the lactation is thus given by:

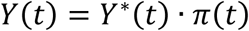

where *t* is the time after parturition in days.

### Unperturbed lactation model

Among the numerous mathematical models developed to study lactation curves, the incomplete Gamma function proposed by Wood [10] has been widely used in different mammals (*e.g.,* rabbit [18], sheep [19], cattle [20]). This model gives a general expression for the dynamics of milk yield along the lactation. In this article, we have selected this model as an example to define the unperturbed lactation curve. Because the structure of PLM is generic, any other lactation model can be used.

The Wood model is given by:

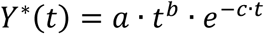

where *Y*^∗^(*t*) is the unperturbed daily milk yield in kg, *t* is the time in days after parturition and *a*, *b*, *c* are positive parameters that determine the shape of the lactation curve (*a* scales the general level of the curve, *b* controls the type and magnitude of the curvature of the function, and *c* regulates the rate of decrease in milk yield after the lactation peak). Values of these parameters can be used to calculate some essential features of the lactation curve such as the time of peak yield (*b*⁄*c*, in days), the lactation persistency, *i.e.*, the extent to which peak yield is maintained (−(*b* + 1) · *ln*(*c*) in kg.d^-1^), or the peak yield (*a* · (*b*⁄*c*)^*b*^ · *e*^−*b*^ in kg) [21].

### Perturbation model

The perturbation model is based on the idea that each single perturbation *i* affecting lactation dynamics can be described as a transient proportional decrease in milk yield, through a sequence of collapse and recovery. Each perturbation can thus be modelled by way of a 3-compartment model (Figure 1) representing the dynamics of the proportion of milk withdrawn from the theoretical unperturbed yield.

**Figure 1.**
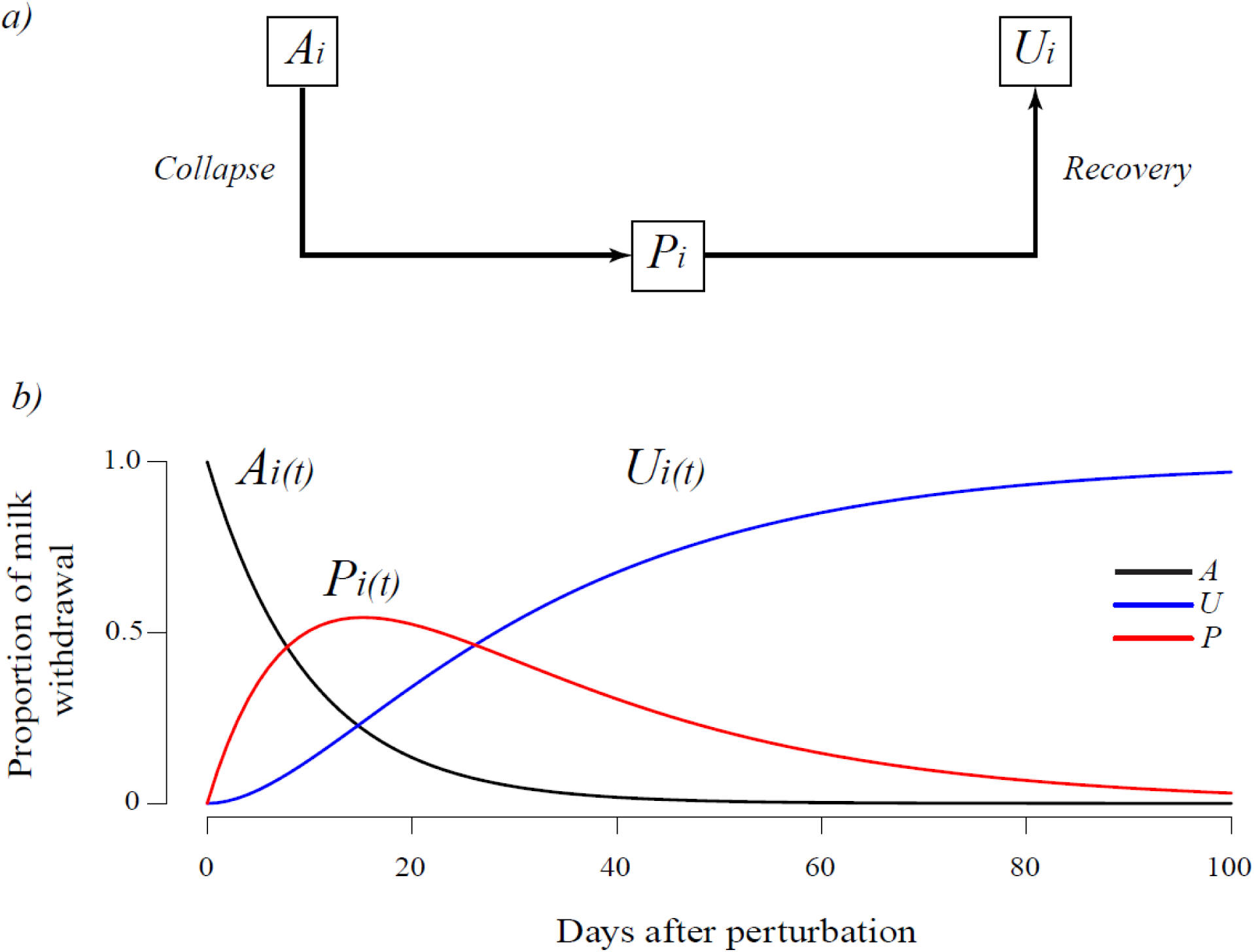
Conceptual model of a single perturbation. *A*: proportion affected by the perturbation, *P*: proportion effectively affected by the perturbation, *U*: proportion unaffected by the perturbation. a) Model diagram and b) Solution dynamics.

The three compartments of the model are: *A*_*i*_, the maximal proportion potentially affected by the *i^th^* perturbation, *U*_*i*_, the proportion unaffected by the *i^th^* perturbation, and *P*_*i*_, the proportion effectively affected by the *i^th^* perturbation. Given the structure of the compartmental model, forming a path from *A*_*i*_ to *U*_*i*_ through *P*_*i*_, and given that the model is defined such as *A*_*i*_ + *P*_*i*_ + *U*_*i*_ = 1, the dynamics of *P*_*i*_ represents the proportional deviation in milk yield.

The perturbation model for a single perturbation *i* is defined by the following simple differential system:

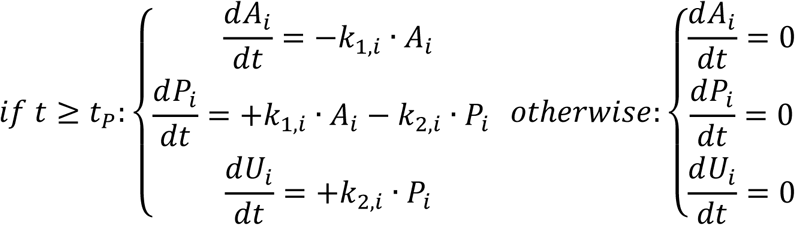

with the following initial conditions at parturition time (*t* = 0):

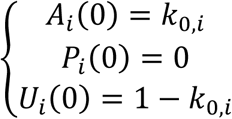

and where *t*_*P_i_*_, is the time of start of the *i^th^* perturbation, *k*_0,*i*_ is the parameter of intensity of the *i^th^* perturbation (*k*_0,*i*_ ∈]0; 1]), *k*_1,*i*_ is the parameter of collapse speed of the *i^th^* perturbation and *k*_2,*i*_ is the parameter of recovery speed of the *i^th^* perturbation.

Assuming that *k*_1,*i*_ ≠ *k*_2,*i*_, the algebraic solution of this differential system is given by:

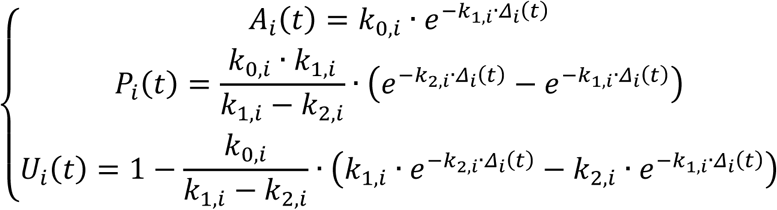

where *Δ*_*i*_(*t*) is the elapsed time since the beginning of the *i^th^* perturbation and is given by:

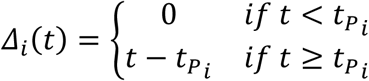

Finally, the perturbation model, including *n* individual perturbations affecting the lactation curve is given by:

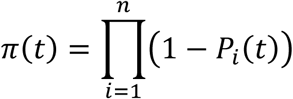

### Model formalism

The detailed algebraic formula of PLM with *n* individual perturbations is given by:

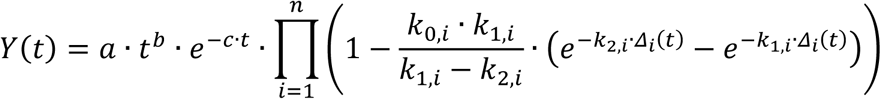

The model includes the three parameters of the Wood model (*a*, *b*, and *c*) to define the unperturbed lactation curve, one parameter to define the number of perturbations affecting the lactation curve (*n*), and four parameters per individual perturbation *i* (*t*_*P_i_*_, *k*_0,*i*_, *k*_1,*i*_, and *k*_2,*i*_) so that the total number of parameters to define PLM is equal to 4 + 4 · *n*.

A simulation of PLM with five perturbations over 300 days of lactation is shown in Figure 2 as an illustration of the model behavior.

**Figure 2.**
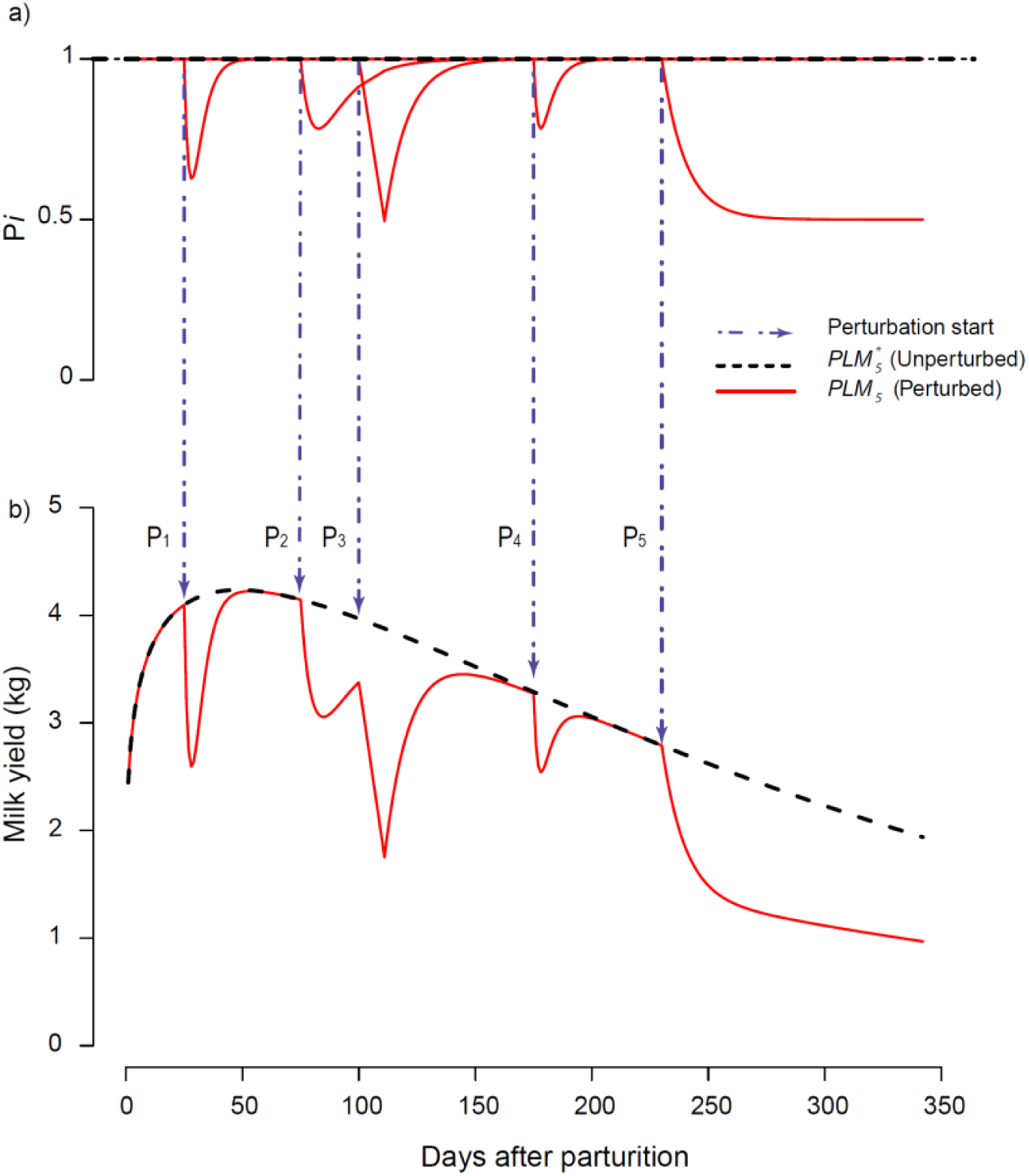
Example of a simulation of the Perturbed Lactation Model (PLM) including five perturbations with a) individual perturbations dynamics expressed as the proportion of unperturbed lactation curve (*Pi*) and b) unperturbed and perturbed milk yield dynamics.

Perturbations were considered individually so that a perturbation can occur within another one (see P_3_ in Figure 2 at *t*_*P*_3__ = 100). Given that individual perturbations are proportional deviations multiplied between them, when a perturbation is added during another perturbation, the new perturbation is a proportion of the already perturbed curve. Moreover, perturbations can be used to simulate the effect of pregnancy (see P_5_ in Figure 2 at *t*_*P*_5__ = 225) with the recovery parameter *k_2,i_* set to zero.

### Fitting procedure

PLM is aimed at detecting perturbations in milk yield time-series data and thus, provide estimates of (1) a theoretical unperturbed lactation curve and (2) the number, timing and shape of the perturbations leading to the observed perturbed lactation curve. A dedicated algorithm was developed in R (R Core Development Team, 2018) with the aim of fitting PLM on lactation data and deriving parameter estimates *a*, *b*, and *c* to characterize the unperturbed lactation curve, *n* to define the number of perturbations and parameter estimates (*t*_*P_i_*_, *k*_0,*i*_, *k*_1,*i*_, and *k*_2,*i*_) for each *i^th^* detected perturbation. Preliminary tests have shown that repeated fittings using different starting values can lead to the detection of perturbations differing in total number and detection order. This raised the question of the theoretical identifiability of the model parameters (for further details on identifiability see [22]) and of the use of a stop criterion to estimate *n*. The structure of the model does not allow a classical identifiability analysis to be performed if *n* is unknown. However, by using the software DAISY (Differential Algebra for Identifiability of Systems [23]), we could assess that for one perturbation the PLM parameters are locally identifiable. To facilitate the identification of the model parameters, we adopted a fitting strategy in two steps: first, performing numerous repeated fittings to estimate the most frequent number of perturbations. In the second step, we fixed as known the number of perturbations detected in step 1 and proceeded to estimate the remaining parameters of the model. This strategy ultimately makes it possible to estimate an optimal number of perturbations and facilitates the estimation of the model parameters.

In the following section, *PLN*_*n*_ stands for PLM with *n* perturbations, *k*_*w_n_*_ stands for the triplet of parameters (*a*, *b*, *c*) of Wood’s model estimated with *n* perturbations (*n* ranging from 0 to *n*_*max*_) and *k*_*P_i,n_*_ stands for the quadruplet (*t*_*P_i_*_, *k*_0,*i*_, *k*_1,*i*_, *k*_2,*i*_) of the *i^th^* perturbation (*n* ranging from 1 to *n*_max_). Since *PLN*_*n*_ combines an estimated unperturbed lactation curve and *n* perturbations, 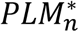 stands for the unperturbed lactation model (*i.e.*, the lactation curve when the n perturbations are removed). *PLN*_0_ (*i.e.*, PLM with zero perturbation) corresponds to the original Wood’s model without any perturbation.

The nls.multstart package (version 1.0.0; [24]) performing non-linear least squares regression with the Levenberg-Marquardt algorithm and with multiple starting values was used for each single fit. Two different sampling schemes of starting parameters were used: random sampling of starting parameters from a uniform distribution within the starting parameter bounds or selection of combinations of starting parameters at equally spaced intervals across each of the starting parameter bounds. These two fitting methods are hereafter referred to as ‘shotgun search’ and ‘gridstart search’ respectively. Starting parameter bounds are defined as follows: *a*: [0; 100]; *b*: [0; 1]; *c*: [0; 1]; *t*_*P_i_*_: [*t*_0_; *t*_3_] (where *t*_0_ and *t*_3_ are the times of first and last records of the dataset); *k*_0,*i*_: [0; 1]; *k*_1,*i*_: [0; 10]; *k*_2,*i*_: [0; 10]. For the ‘shotgun search’, the number of random combinations of starting parameters was set to 100,000. For the ‘gridstart search’, the number of combinations of starting parameters (*i.e.*, the size of the grid), was set to five for parameters *a*, *b*, *c*, *k*_0,*i*_, *k*_1,*i*_, *k*_2,*i*_ and to 10 for the parameter *t*_*P_i_*_. Consequently, for the fit of one perturbation (*i.e.*, estimating 3 + 4 = 7 parameters) the number of tested combinations of starting parameters was 7^6^ x 10 = 1,176,490. For both search methods, the best model was selected on the basis of the lowest Akaike Information Criterion (AIC) score [25].

The whole fitting procedure includes repetitions of a fitting sequence that proceeds by successive addition of perturbations. This fitting sequence is defined in such a way that the estimate of the parameters of each new perturbation is obtained while the parameters of the previously added perturbations are kept fixed. Therefore, the fitting of *PLN*_*i*_ provides parameters estimates for the new added *i^th^* perturbation and for a new version of Wood model’s parameters *k*_*w_i_*_ (*i.e.*, each time a new perturbation is added, a new version of the unperturbed lactation is refined). For a given lactation dataset composed of daily milk yield records, the preliminary fitting of *PLN*_0_ (*i.e.,* the original Wood’s model without any perturbation) was first performed to estimate *k*_*w*0_. Then, the fitting sequence starts by the fitting of *PLN*_1_ (*i.e.,* PLM with 1 perturbation) thus providing estimates *k*_*w*_1__ and *k*_*P*_1,1__. Then, the fitting of *PLN*_2_ consists in estimating *k*_*w*_2__ and *k*_*P*_2,2__ with *k*_*P*_1,2__ fixed equal to *k*_*P*_1,1__. Then, the fitting of *PLN*_3_ consists in estimating *k*_*w*_3__ and *k*_*P*_3,3__ with *k*_*P*_1,3__ and *k*_*P*_2,3__ fixed equal to *k*_*P*_1,2__ and *k*_*P*_2,2__, respectively. The procedure is applied stepwise until the maximum number of perturbation *n_max_* is reached. This maximum number is an *a priori* user defined value to fix a stop criterion. Preliminary tests have shown that setting *n_max_* = 15 was sufficient. The end of the fitting sequence consists in reordering the *n_max_* detected perturbations in decreasing order according to the time of perturbation *t*_*P_i_*_ (the original obtained order of perturbations is based on the opportunities found by the fitting procedure to improve the goodness-of-fit for each added perturbation).

Finally, the whole fitting procedure is carried out following the 3 following steps:

Step1: Repeat 100 times the fitting sequence with the ‘shotgun search’ and *n_max_* = 15.

Step2: Compare the fitting results of the 100 repetitions obtained in Step1 and identify perturbations systematically detected at *t*_*P_i_*_ ± 3 days. This was performed by counting, for the 15 perturbations over the 100 fitting results, the number of occurrences of the rounded value 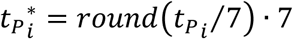. Step1 provides the optimal number of perturbations denoted *N* (*i.e.,* the value of n giving the best fit) with an estimate of *t*_*P_i_*_ for each perturbation (calculated as the median of the *t*_*P_i_*_ with the same rounded value 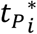).

Step3: Perform the fitting sequence with the ‘gridstart search’, with *n_max_* = *N* and with starting parameters bounds for each *t*_*P_i_*_ reset to [*t*_*P_i_*_ − 10; *t*_*P_i_*_ + 10]. This last fit provides the final estimates *k*_*w_N_*_ and (*k*_*P*_1,*N*__, …, and *k*_*P_N,N_*_) characterizing respectively the best fit for the unperturbed model and the *N* detected perturbations. The Root Mean Square Error (RMSE) was calculated to indicate the goodness-of-fit of *PLN*_*N*_. Additionally, the percentage of loss ′*L*′ was calculated using the formula 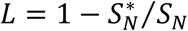 where 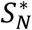 and *S*_*N*_ are respectively the total milk yield over [*t*_0_; *t*_3_] calculated with 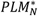 (the unperturbed curve corrected from N perturbations) and *PLN*_*N*_ (the perturbed curve with N perturbations).

To provide complementary information on lactation time-series and refine PLM outputs analysis, the model of Grossman et al. [26] was also fit to lactation data as described in Martin and Sauvant [27]. This fitting cuts the lactation period into three stages corresponding to early, middle and late stages (respectively intervals [*t*_0_; *t*_1_]: increasing phase, [*t*_1_; *t*_2_]: plateau-like phase, and [*t*_2_; *t*_3_]: decreasing phase). This triphasic model, based on a smoothing logistic transition between intersecting straight lines, specifies the cut points of the three stages (instead of *a priori* number of days in milk). This fit was performed using the ‘gridstart search’ with [*t*_0_; *t*_3_] as starting parameters bounds for the interval terminals *t*_1_and *t*_2_.

### Dairy goat dataset

In this study we used data from 181 goats (94 Alpine and 87 Saanen) born between 2009 and 2017. Data concerned 319 lactations (126 primiparous and 193 multiparous; parity ranging from 1 to 7) including 80,773 milk records from the dairy goat herd of the INRA-AgroParisTech Systemic Modelling Applied to Ruminants research unit (Paris, France) between 2015 and 2018. Records are shown in supplementary Figure 1 by breed and parity. All lactations considered had at least one record in the first 5 days of lactation and a last record between 150 and 358 days of lactation (no extended lactation included).

### Statistical analysis

All statistical analyses were performed using R (R Core Development Team, 2018). Fixed effects of breed (Saanen *vs.* Alpine) and parity (1 *vs.* 2 and more) were tested on parameters of Wood, with and without the changes made from PLM model. It was also tested on estimated peak milk yield, peak time, total milk yield over [t_0_; t_3_], the number of perturbation and the rate milk loss using a mixed analysis of variance model with goat as a random factor. Fixed effect of lactation stage (early *vs.* middle *vs.* late) was tested on RMSE and on PLM parameters *t*_*P*_, *k*_0_, *k*_1_, *k*_2_ with a mixed analysis of variance model with parity as a random factor. Pearson linear correlations were calculated for PLM parameters: intra-class of breed and parity for *a*, *b*, *c*, *N*, and *L* and intra-class of stage of lactation for *t*_*P*_, *k*_0_, *k*_1_, and *k*_2_.

## RESULTS

Lactation duration ranged from *t*_0_ = 1.2 ± 0.6 to *t*_3_ = 270.3 ± 40.8 days in milk. Early, middle, and late lactation stages determined with Grossman’s model were 1.2 to 34.4, 34.4 to 171.0, and 171.0 to 270.3 days, respectively.

### Fitting

The fitting procedure converged for the 319 lactations and detected a total of 2,354 perturbations with an average of 7.4 perturbations per animal per lactation. Figure 3 shows the fitting of PLM on one lactation dataset. The fitting results on individual lactations exhibiting the minimum and maximum values for the RMSE (0.1 kg and 0.4 kg) are provided in supplementary Figure 2. The number of perturbations varied between 4 and 11, the percentage of milk loss between 2% and 19%, the total unperturbed milk yield was between 393 kg and 1,557 kg and the record interval length was between 1 and 5 days for *t_0_* and between 165 and 358 days for *t_3_*. During the first fitting steps, the Wood’s parameters were stabilized on average after the detection of the first 4 perturbations (supplementary Figure 3). This indicates the robustness of the unperturbed curve.

**Figure 3.**
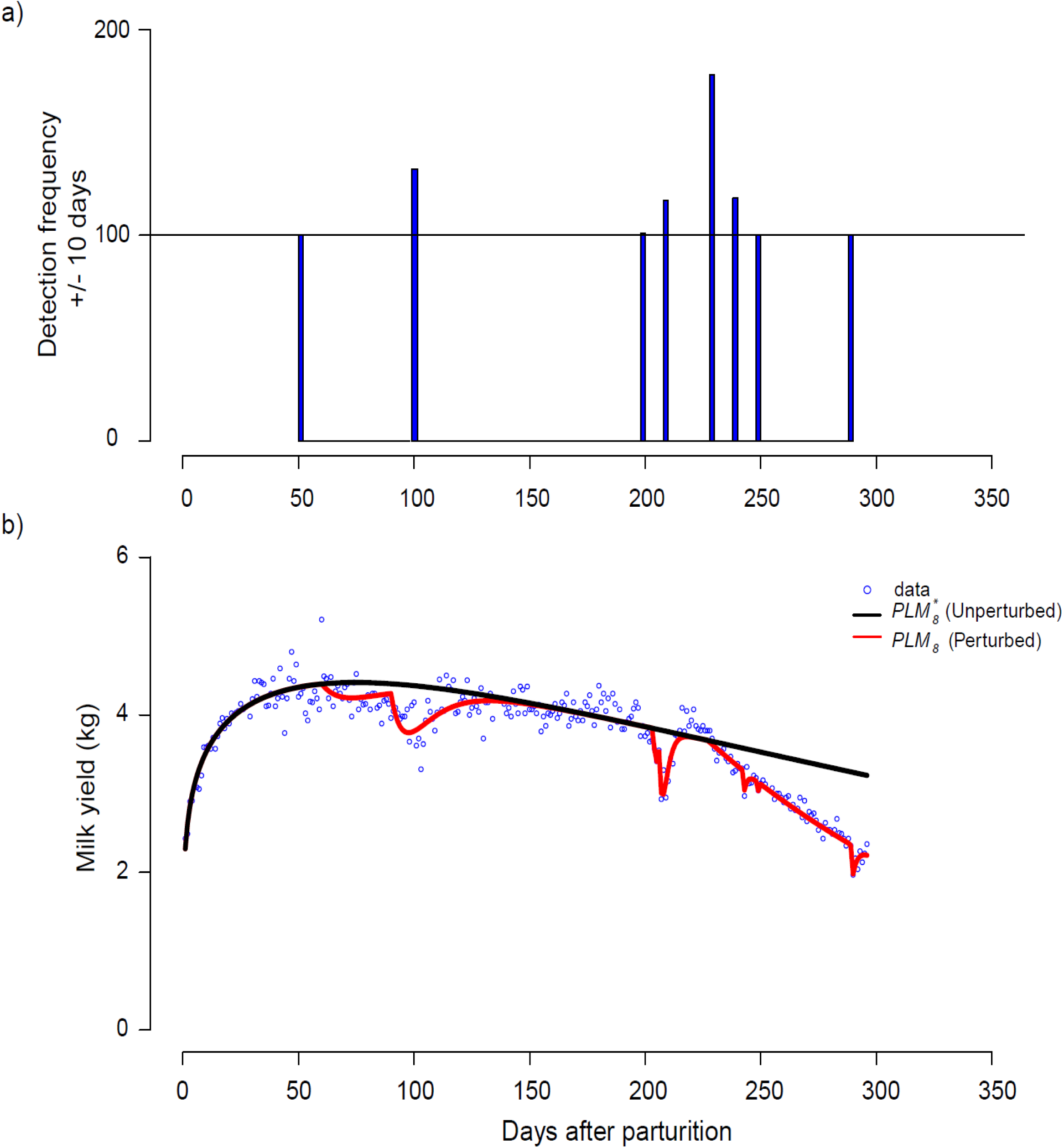
Example of the perturbed lactation model fitting procedure result on a one goat lactation dataset. a) frequency of detection of a single perturbation within +/- 10 days; b: unperturbed and perturbed lactation models plotted against data.

Descriptive statistics of the results obtained from the fitting procedure of *PLN*_*N*_ are given in Table 2 by breed and parity and are compared to the results obtained with *PLN*_0_, corresponding to an adjustment of the Wood model without any perturbations. The value for the parameter *a* greatly increased between the Wood model and *PLN*_*N*_. The values for parameters *b* and *c* decreased between the Wood model and *PLN*_*N*_. As a consequence, values for peak milk and time of peak increased between the Wood model and *PLN*_*N*_. Both models did not give a similar level of variance of error according to breed or parity level. Regarding the quality of fitting, the RMSE values showed a fairly significant decline between the Wood model (0.4 ± 0.1 kg) and *PLN*_*N*_ (0.2 ± 0.1 kg). Considering explicit perturbations in the fitting of the Wood model with PLM compare to fitting directly the Wood function to data led to a decrease in RMSE, reflecting an improvement in the goodness of fit.

**Table 2.**
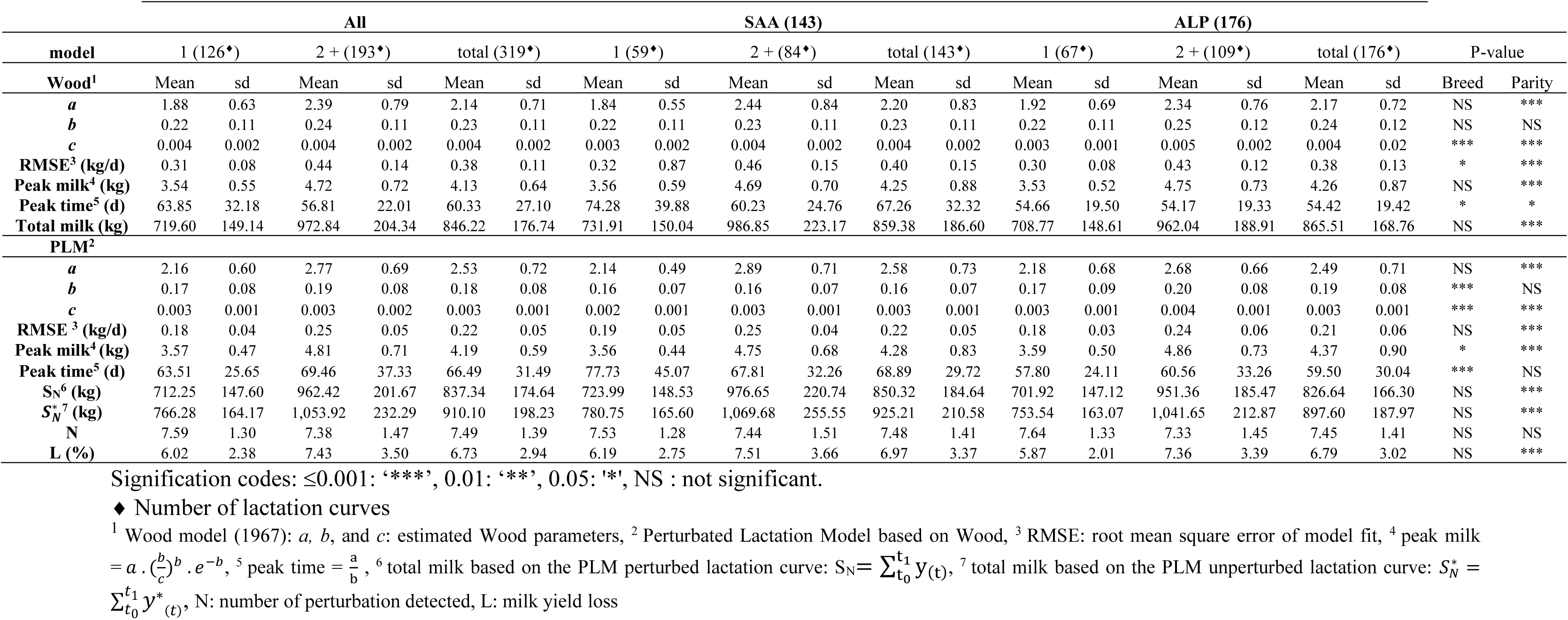
Results of the fitting procedure: comparison between breeds and lactation numbers across the models and variables.

### Unperturbed lactation curve

Descriptive statistics of the parameters *a*, *b* and *c* for the unperturbed lactation curves (for both models: 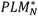 and Wood model) are presented in Table 2 for the overall dataset, breed and parity. The parameter *a*, which drives the general scaling of the curve, was not significantly different for the two breeds (Alpine: 2.49 ± 0.71; Saanen: 2.58 ± 0.73). Consequently, no significant breed effect was found for the peak milk or for the total unperturbed milk production. The same statistical effects were found with the Wood adjustment without perturbation. The parameter *a* was significantly affected by the parity of the lactation, with first lactations having a lower value for parameter *a* than the two and more parities (Table 2). Consequently, there was a significant parity effect on the peak milk and on the total milk production. The parameter *b*, which drives the curvature of the lactation curve, was significantly affected by breed. Alpine goats exhibited higher values of *b* compared to Saanen goats (Alpine: 0.19 ± 0.08; Saanen: 0.16 ± 0.07). Parity also had a significant effect on the parameter *b*, with first lactations having a lower value for parameter *b* than two and more lactations. Regarding the parameter *c*, which drives the rate of decrease of milk production after the peak, both parity and breed effects were highly significant. Alpine goats exhibited a same value for the parameter *c* than the Saanen goats (Alpine: 0.003 ± 0.001; Saanen: 0.003 ± 0.001). For this parameter, first lactations had a lower value than two and more lactations (Primiparous: 0.002 ± 0.001; Multiparous: 0.003 ± 0.001). The peak time of the unperturbed curve, resulting from both *b* and *c* parameters, was significantly affected by breed, with Saanen goats exhibiting a peak 14 days later in lactation than the Alpine goats. The statistical effects found for 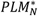 parameters were consistent with the effects found for the Wood model (*PLN*_0_), except for the peak time. Regarding peak time, the Wood model peak time was slightly affected by both breed and parity, while for the 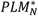 peak time, breed had a very significant effect and parity was not significant.

Individual unperturbed lactation curves obtained with 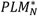 for increasing parities are shown in Figure 4. Some of these individual adjusted curves were considered as atypical, in the sense they departed from the general shape of the Wood model. An individual lactation was considered “atypical” if the persistence estimated by PLM, *i.e.* the value of parameter *c*, was an outlier, defined as a value either 3 times above the inter-quartile range (IQR) (above the third quartile of the distribution for the *c* parameter) or 3 times below the IQR (below the first quartile of the distribution for the *c* parameter). However, it is important to note that atypical curves observed in the dataset were biologically true. A total of 18 out of the 319 analyzed curves were classified as atypical. Generally, these atypical curves come from the same goat in different parities or for primiparous that have not started the second parity. Peaks of milk of the unperturbed lactation curve were on average increased by 27.47% between the first parity and the second parity, by 9.46% between the second parity and the third parity, and by −0.29% between the third parity and the fourth parity (Figure 4). The total milk production for the unperturbed curve was increased by 32.55% between the first parity and the second parity, 5.20% between the second parity and the third parity, and by 1.01% between the third parity and the fourth parity. These results are consistent with Arnal et al. [28].

**Figure 4.**
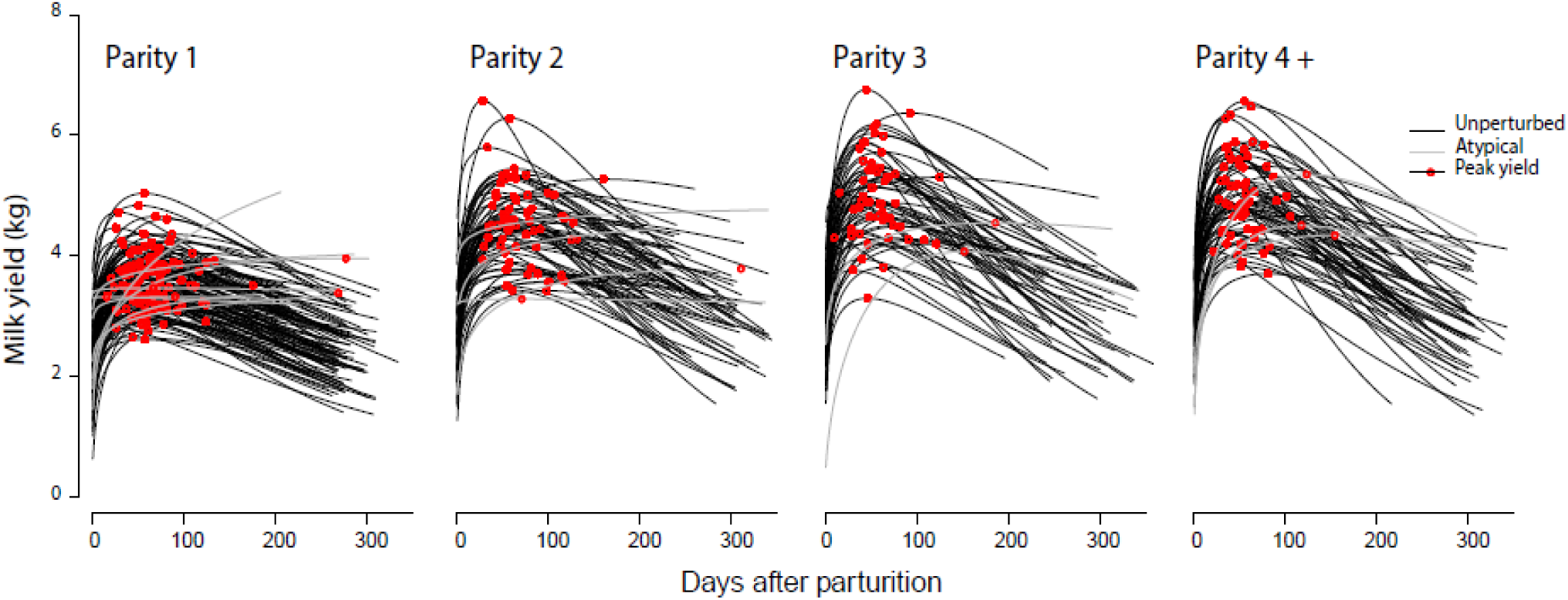
Individual unperturbed curves extracted from data after removal of the estimated perturbations using PLM for increasing parity number (fit on 319 lactation data; atypical curves correspond to outlying estimates of the parameter *c* governing milk persistency).

The Pearson linear correlation matrix by breed and parity between parameters of 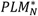 is shown in Figure 5 (panels a and b). A strong negative correlation was found between *a* and *b* (−0.65), indicating that high values of *a* (scaling of the lactation curve), were associated with low values of *b* (shaping the curve). A positive correlation was found between the parameters *c* and *b* (0.64) indicating a positive association between the shape of the curve and the rate of decrease of lactation. Finally, a low negative correlation between *c* and *a* (−0.11) was found. These results are consistent with the well-known features of lactation curves: higher milk at peak yield being associated with higher speed of decline after peak. Several factors (*e.g.* breed, parity, seasonality, and season of kidding) can affect characteristics of the lactation curve. The differences found in this study between primiparous and multiparous goats are consistent with previous studies [28, 29] with primiparous goats being less productive, with a lower peak yield and a greater persistency. Despite the lack of a significant effect of parity, our results are consistent with previous studies [29] where primiparous goats had a peak later than multiparous (see Table 2). The strong breed effect we observed on peak time is consistent with previous studies [29] with Saanen goats having a peak yield later than Alpine goats.

**Figure 5.**
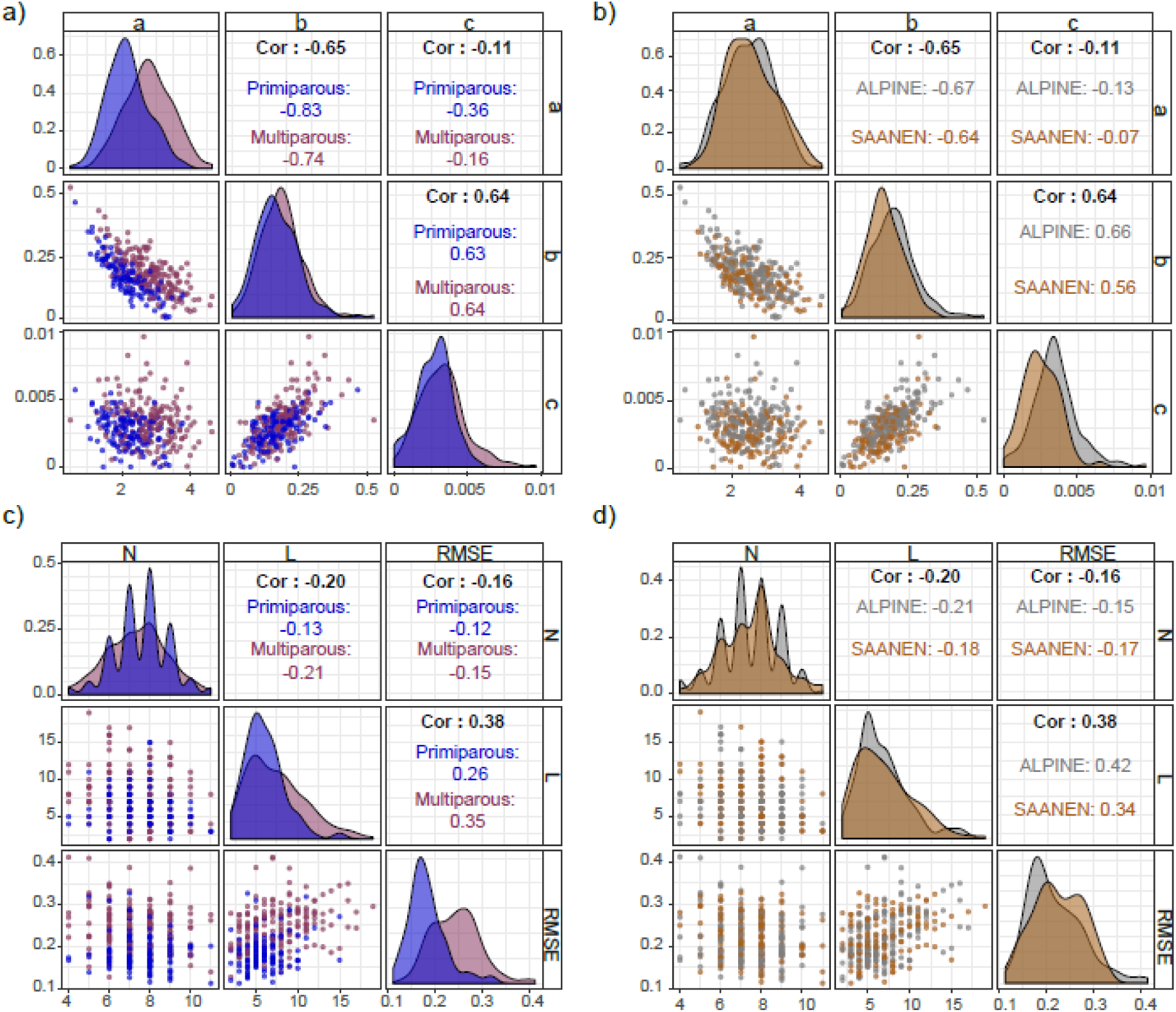
Pearson linear correlation matrix of PLM parameters estimates: panels (a) and (b): the *a, b, c* parameters defining the unperturbed curve (a: by parity and b: by breed). Panels (c) and (d): the number of perturbations N, milk loss and RMSE (c: by parity and d: by breed).

### Number of perturbations and milk loss

The effects of parity and breed on the total number of perturbations were not significant. Total number of perturbations was 7.59 for the primiparous, 7.38 for the multiparous, 7.45 for the Alpine and 7.47 for the Saanen. By contrast, the rate of milk yield loss after perturbation was significantly affected by the parity. A Pearson linear correlation matrix by breed and parity between 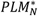 estimates for the number of perturbations (*N*), percentage loss of milk yield (*L*), and goodness of fit RMSE was also carried out (Figure 5, panels c and d). A positive correlation was noted between RMSE and milk loss (0.38). However, weak negative correlations between the number of detected perturbations and RMSE (−0.16), and the number of perturbations and the milk loss (−0.20) were also noted. Distributions of *N*, *L* and RMSE showed an even larger difference according to the parity than to the breeds. These results show that it is not the number of perturbations that contribute the most to the loss in milk yield over the lactation.

### Perturbation timing and shape

Table 3 gives descriptive statistics on the parameters of PLM characterizing the 2,354 perturbations detected during the fitting procedure: time *t*_*P*_, intensity *k*_0_, collapse speed *k*_1_ and recovery speed *k*_2_ according to the lactation stage determined with Grossman’s model. Most of the perturbations were detected during the late stage of lactation (n = 1,063). The number of perturbations tended to decrease in middle stage (n = 1,054) and for early stage (n = 237). The parameter *k*_0_ increased from early, middle and late lactation stage (Table 3). These results suggest that throughout the lactation process, perturbations become more intense. The parameter *k*_1_ decreased from early to late stages of lactation. This suggests that perturbations tended to be sharper at the beginning of lactation, with a high speed of collapse and recovery, while they tended to be smoother than the lactation progressed

**Table 3.** Descriptive statistics of perturbation parameters for the 2,354 perturbations detected by the perturbed lactation model in the dairy goat lactation dataset.

|  | Stage of lactation (2,354) |  |  |  |  |  |
| --- | --- | --- | --- | --- | --- | --- |
|  | Early (237) |  | Middle(1,054) |  | Late (1,063) |  |
| Perturbations | Mean | Sd | Mean | sd | Mean | sd |
| $tp$ : time | 33.8 | 34.0 | 107 | 63.0 | 202 | 60.0 |
| $k_0$: intensity | 0.450 | 0.331 | 0.506 | 0.349 | 0.672 | 0.359 |
| $k_1$ : collapse | 4.01 | 4.17 | 3.41 | 3.87 | 2.76 | 3.69 |
| $k_2$ : recovery | 1.13 | 1.96 | 1.18 | 1.79 | 0.95 | 1.71 |

The PLM parameter *k*_0_, which drives the intensity of the perturbation, varied considerably between 0.001 and 1 (set as a boundary). The parameter *k*_1_ (which drives the collapse speed of the perturbation), and the parameter *k*_2_ (which drives the speed of recovery) varied between 0 and 10. A gradient according to the stage lactation was noted for these parameters. A gradual increase in *k*_0_ and a gradual decrease in *k*_1_ and *k*_2_ according to early, middle and late lactation stages was noted (Table 3). In the late stage, 30.20% of the perturbations were detected with a parameter *k*_2_ equal to 0, which implied a perturbation without any recovery period. Among these perturbations, 85.39% had a *k*_0_ value equal to 1. On the other hand, in the early and middle stages, the perturbations detected with an *k*_2_ equal to 0 were 1.70% and 7.07%, respectively.

## DISCUSSION

### Combining two types of models

In this study, we described the PLM model proposed as a tool for extracting simultaneously perturbed and unperturbed lactation curves from daily milk time-series. The key original feature of PLM is to combine an explicit representation of perturbations with a mathematical representation of the lactation curve.

Regarding the mathematical representation of the lactation curve, the structure of PLM is generic and any equation can be used to describe the general pattern of milk production throughout lactation (see appendix including Figure 4 showing illustration of results with other lactation models). The Wood model [10] was chosen in this study as it is one of the most well-known and commonly used mathematical model of lactation curve. Behind the choice of considering a general pattern of lactation that is distorted by perturbations, the biological assumption is that the dairy female has a theoretical production potential (the unperturbed curve) corresponding to the expression of its genetics in a given environment. This genetic potential may not be fully expressed in the farm environment because of perturbations (the perturbed curve).

Regarding the representation of perturbations, we chose an explicit formalism with a compartmental structure for every single perturbation. With this conceptual choice, PLM overcomes limitations of recent models developed for capturing perturbations [15–17]. It allows the capture of multiple perturbations with contrasted features: from a sharp and short drop (for instance due to a diarrhea episode) to a long and slow decrease (for instance due the gestation status). PLM also allows to determine the time at which the perturbations occur during the lactation. This last point is of great interest to add value to on-farm data where challenges imposed to animals do not result from controlled trials and arise from the farm environment.

By combining a general model of lactation curve with an explicit model of perturbations, PLM provides two key outputs: first, the unperturbed curve of the lactating female reflects its production potential in a non-perturbed environment, and second the perturbed curve which reflects the production permitted by the farm environment. The PLM parameters (*k*_0,*i*_, *k*_1,*i*_ and *k*_2,*i*_) provides the most useful information on characteristics of the perturbed lactation curve including scale and shape for each perturbation. Indeed, by providing a perturbed curve, we give an estimate of the number of perturbations and for each perturbation an estimate of its time of start *t*_*P*,*i*_, intensity *k*_0,*i*_, collapse speed *k*_1,*i*_ and recovery speed *k*_2,*i*_. This not only allows PLM to be flexible in capturing different types of perturbations (*e.g.,* gestation, drying, disease), but also to produce metrics to compare the effect of these perturbations on milk yield. In such cases, and by introducing the information concerning these perturbations as an explicit component in the Wood model, we force the model to take into account these perturbations to build the unperturbed curve.

With the development of on-farm technology measurements, an interesting perspective for PLM is to be used to assess other biological time-series data, such as body weight changes, dry matter intake, and hormones dynamics during lactation.

### Fitting algorithm

Beyond the original concepts behind PLM, a key methodological development has been the fitting algorithm. The number of parameters to be determined is substantially important, including the Wood parameters of the unperturbed curve (3 parameters), and PLM parameters (4 parameters for each perturbation). To overcome the difficulty of estimating a high number of parameters, a 2-step algorithm was implemented. The first step of the procedure was to determine Wood parameters and the time when the perturbation starts. The second step of the procedure was to determine PLM parameters. Another difficulty concerned the choice of a maximum number of perturbations. After several attempts, this 2-step algorithm was selected for three main reasons. The first one was related to the visual quality of the fitting results itself. Indeed, the obtained fitted curve is always very close to what would have been drawn after simply looking at the raw data and wondering what the lactation curve would be without perturbations. This proximity to what could have been inferred was considered decisive, yet subjective. The second reason was related to the issue of finding the number of perturbations. The PLM procedure allows an automated determination of an optimal number of perturbations, without *a priori* estimates or use of an arbitrarily chosen stopping criterion. Preliminary results have showed that allowing a maximal number of 15 perturbations to be detected in the first step of the algorithm was enough for the considered dataset. The third reason pertained to the model parameters identifiability issue [22]. Since the fitting is based on a huge number of repeated fittings from which the systematically detected times of perturbations are retained, the 2-step fitting algorithm facilitates the practical identifiability of the model parameters. Indeed, the overall fitting algorithm was applied several times to the same dataset. Given that obtained parameter estimates were the same between the different runs, not only it strengthens the convergence properties of the algorithm but also it guarantees model parameters identifiability.

Fitting results (see Figure 6) have shown that, in some cases, parameter estimates characterizing an individual perturbation reached their initial upper boundaries (1 for parameter *k*_0,*i*_ and 10 for parameters *k*_1,*i*_ and *k*_2,*i*_). This situation concerns perturbations with a narrow and deep peak-shape. By construction, the value of the parameter *k0* (intensity of the perturbation) is a proportion and thus not supposed to exceed 1. For the parameters *k*_1_ and *k*_2_, a value of 10 already represents a very abrupt collapse or recovery, respectively. These results are therefore considered relevant. However, a next step may be to test the model on a larger dataset to assess the need to broaden these boundaries. Furthermore, another working step will consist in developing an application where the settings of the PLM algorithm can be user-defined. For instance, the maximal number of detectable perturbations, the size of the search grid in step one, or boundaries of parameters.

**Figure 6.**
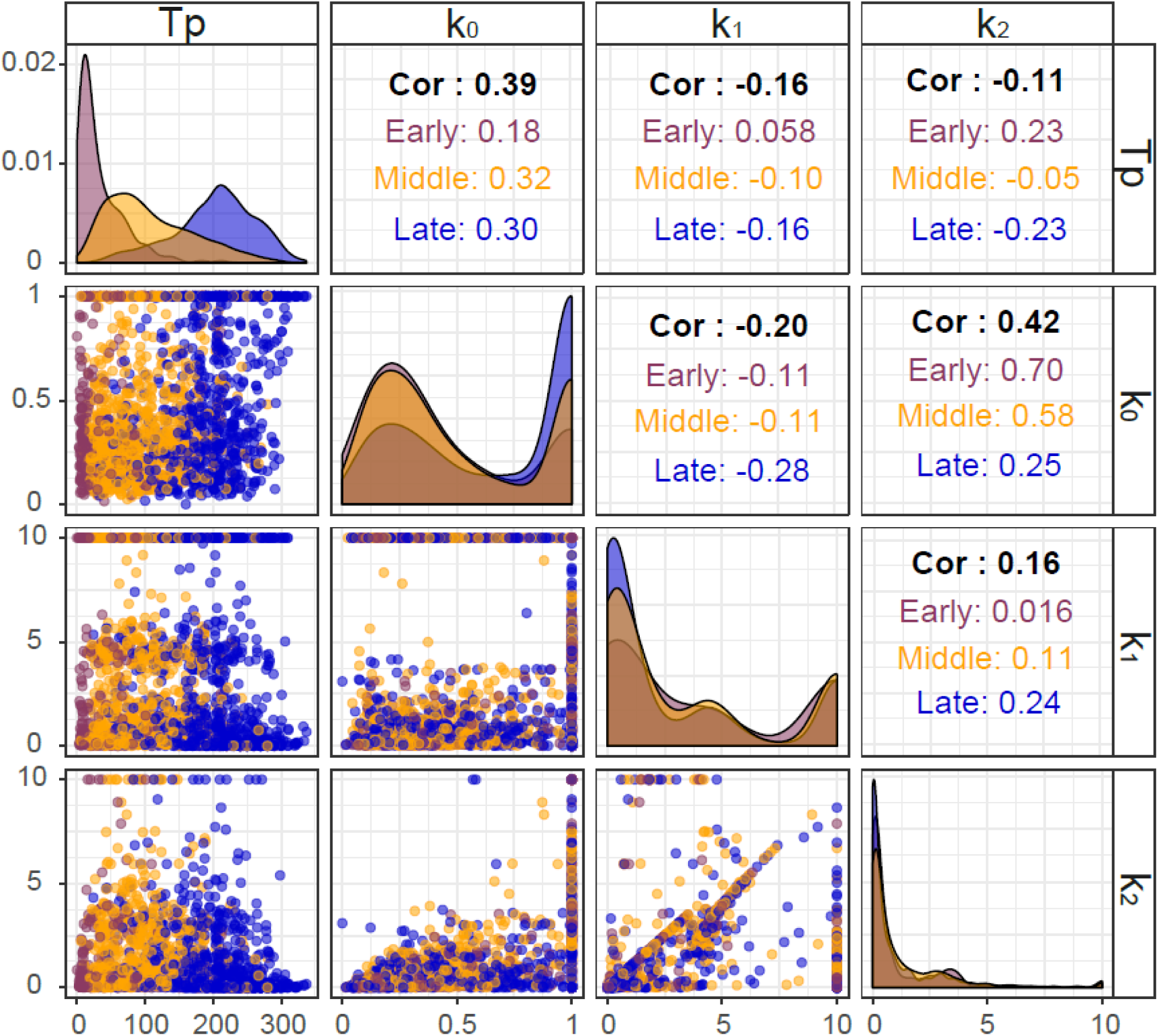
Pearson linear correlation matrix on the PLM parameters by stage of lactation: tp: perturbations times detected; *k0*: intenstity, *k1*: collapse and *k2*: recovery of perturbation.

### Phenotyping tool

PLM was developed to improve the ability to phenotype animals by extracting biological meaningful information from raw data. The unperturbed curve fitted by PLM makes it possible to compare animals based on their potential of milk production. With this information, animals can be ranked based on the production level they would have achieved in a non-perturbed environment, instead of being ranked based on the measured production level assuming no perturbations were encountered. This ranking may be of interest for the famer’s breeding strategy, avoiding the culling of animals that have faced a challenge decreasing their production milk while still having high genetic merit.

The perturbed curve and the characteristics of each perturbation (time, intensity, collapse and recovery) open the perspective of working on perturbations as such and using this information for breeding and management. As a phenotyping tool, PLM can be useful for genetic selection. Studying characteristics of perturbations throughout many lactations of a large number of individuals and linking them to genetic or genomic information opens perspectives to evaluate their heritability and their potential genetic impact. PLM can also be a valuable tool for on-farm management. Linking perturbations with other information on the animals, such as lactation stage, parity, gestation stage, can help to detect sensitive periods where perturbations are more likely to occur. By cross-checking information on perturbations from all animals with information on the farm environment (for instance temperature, feed availability), it would be possible to detect synchronous occurrences of perturbations and link them to farm environment or management practices during times of stress. With this better understanding of environmental effects on animal production, preventive measures on the farm could be made.

Understanding the effects of the environment on farm animals and understanding how they cope with perturbations during crucial times could help to gain insights on resilience and robustness. These complex dynamic properties are highly desirable to face the changes occurring in the livestock sector [30]. While the conceptual framework to work on resilience and robustness is now well defined in animal sciences, we still need operational metrics [31]. Such metrics have been proposed for a single perturbation by Revilla et al., [15] and Sadoul et al. [17]. Taking into account this type of information can provide a proxy to estimate the frequency and severity of disorders such as clinical mastitis [32]. Studying perturbations in lactation curves also makes it possible to compare animals facing the same stress and detect the ones with the greatest adaptive capacities. Finally, the on-farm early detection of perturbations in milk yield can provide farmers with an alert system on udder health. Recently, Huybrechtset al. [33] tested and developed the synergistic control concept for early detection of milk abnormalities in dairy cows based on detection of shifts in milk yield per hour. Of the 49 mastitis cases, 31 cases were detected using this methodology at the same time or earlier than they were detected by the farmer.

To our knowledge, existing metrics for dropped milk yields per day in the lactation curve, as proposed by Elgersma et al. [11], are based on a variance approach applied to the whole curve. Fluctuations in milk yield are summarized with a single statistical measure. Complementary to this type of approach, PLM can decompose the whole curve and characterize each perturbation, with metrics that are consistent with the concept of resilience of each and subsequent perturbation. The PLM model offers a way of quantifying the consequences of external factors and exploring hypotheses about the biological types of responses due to specific perturbations. By giving a biological meaning to these parameters, we can reconcile a phenotyping tool with the opportunity of an explanatory selection approach.

A major limitation of PLM results in its dependency to the quality of data. Indeed, if data are recorded with a low accuracy (due to technical problems of measurements), the outputs of PLM do not have consistency as detected perturbations have nothing to do with perturbations of the lactation curve, but are related to accuracy problem. In addition, PLM has been developed with daily records. It will be necessary to evaluate if PLM can operate correctly with less frequent data. Finally, PLM is based on the concept of a theoretical unperturbed curve of milk production, considered as a potential, and used to determine deviations that reflect perturbations. This rationale for an underlying potential is debatable from a biological point of view. Nevertheless, from a strict mathematical point of view, we considered this approach as valuable to provide a tool for interpreting data. The application of PLM on other datasets from other species could provide information to further evaluate this point.

### Conclusion

By combining a general description of the lactation curve with an explicit representation of perturbations, the PLM model allows the characterization of the potential effects on milk production, allowing to assess animal genetics, and the deviations induced by the environment, reflecting how animals cope with real farm conditions. The translation of raw time series data into quantitative indicators makes it possible to compare animals’ phenotypic potential and bring insights on their resilience to external factors. In that sense, PLM could be used as a valuable phenotyping tool and it contributes to provide decision solutions for dairy production that are grounded in a biologically meaningful framework. Further modelling studies should strive for integrating high throughput data analysis with such biological framework.

## Acknowledgments

We gratefully acknowledge the team at the INRA UMR 791 Modélisation Systémique Appliquée aux Ruminants (Paris, France) experimental installation for the care of the animals and their work to provide robust performance data. Special thanks to Dr. R. Muñoz-Tamayo for his conscientious reading and meticulous corrections of the manuscript. This work was carried out with the financial support of the ANR-Agence Nationale de la Recherche – The French National Research Agency under the “Deffilait project” (ANR; project: ANR-15-CE20-0014). This preprint has been reviewed and recommended by Peer Community In Animal Science (https://doi.org/10.24072/pci.animsci.100001).

