## Supporting Figures for "Lactation curve model with explicit representation of perturbations as a phenotyping tool for dairy livestock precision farming"

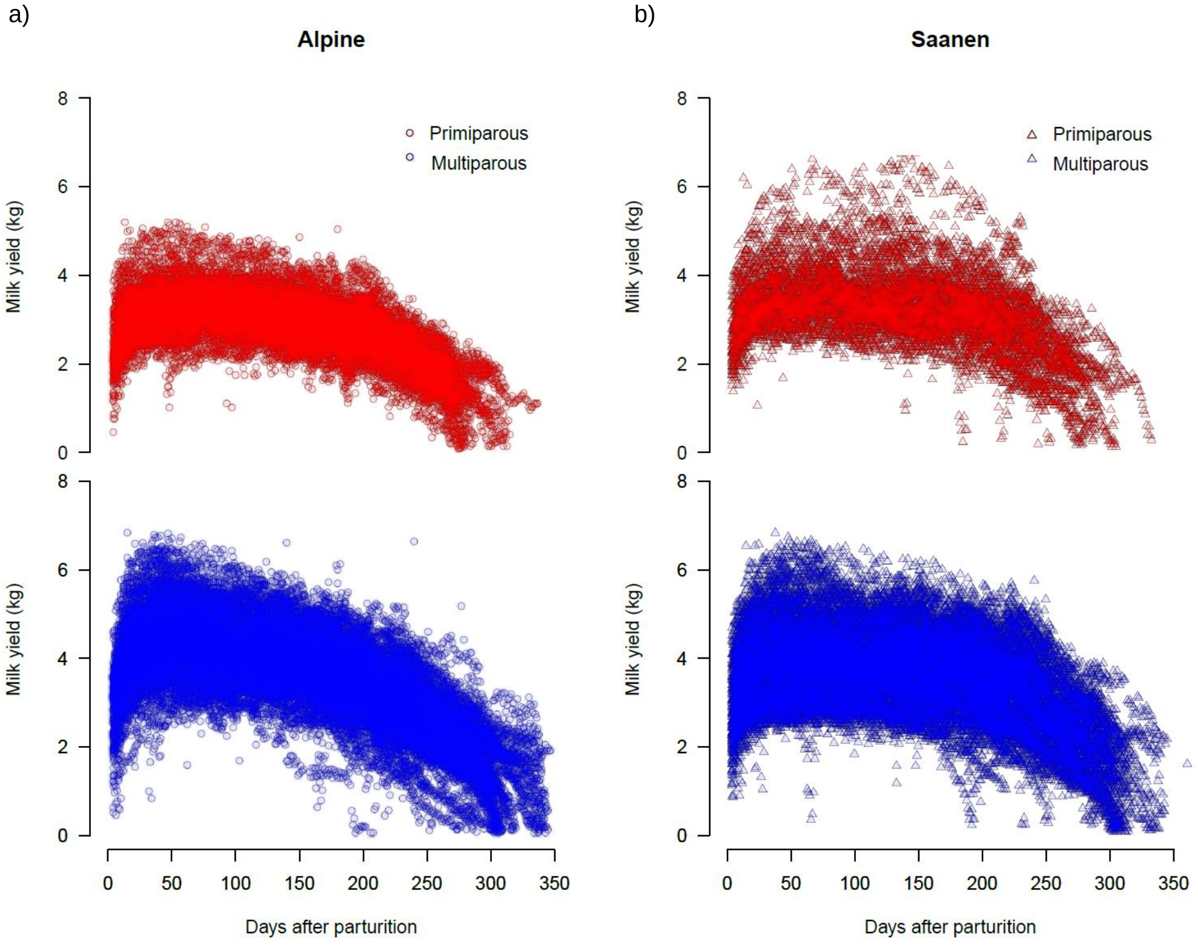


S1. Dairy goat data set: daily milk time series data over lactation (n=319 goats, 80,773 records) by breed (a: Alpine and b: Sannen) and parity (primiparous in red and multiparous in blue).

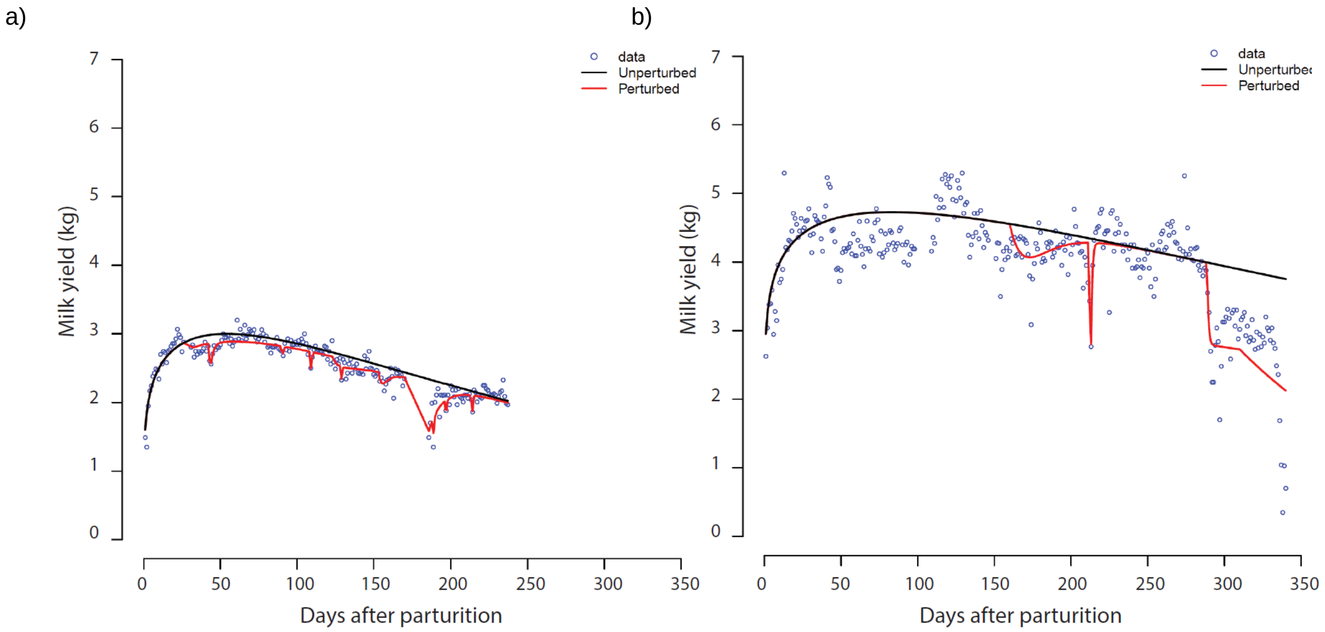


S2. Example of the perturbed lactation model fitting procedure result on a lactation dataset. a) The fitting results on individual lactations exhibiting the minimum values for the RMSE (0.11kg) ; b: The fitting results on individual lactations exhibiting the maximum values for the RMSE (0.41kg);

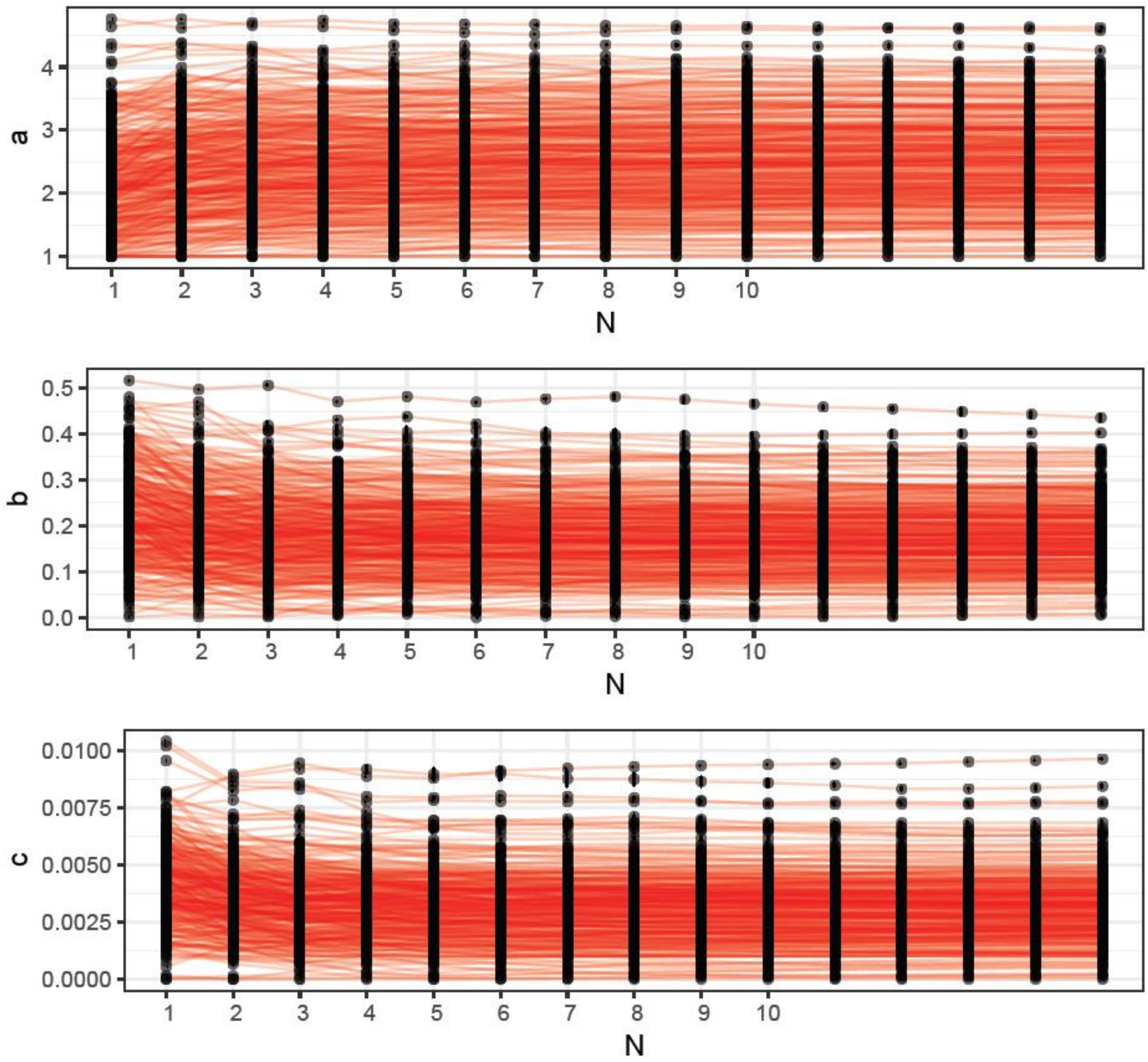


S3. Evolution of the Wood's parameters (a, b and c) during the detection of the 15 perturbations of the PLM first step for the 319 goats studied. Each red line represent the result mean of the repeat of the 100 times fitting sequence with the‘shotgun search’ and Nmax=15.

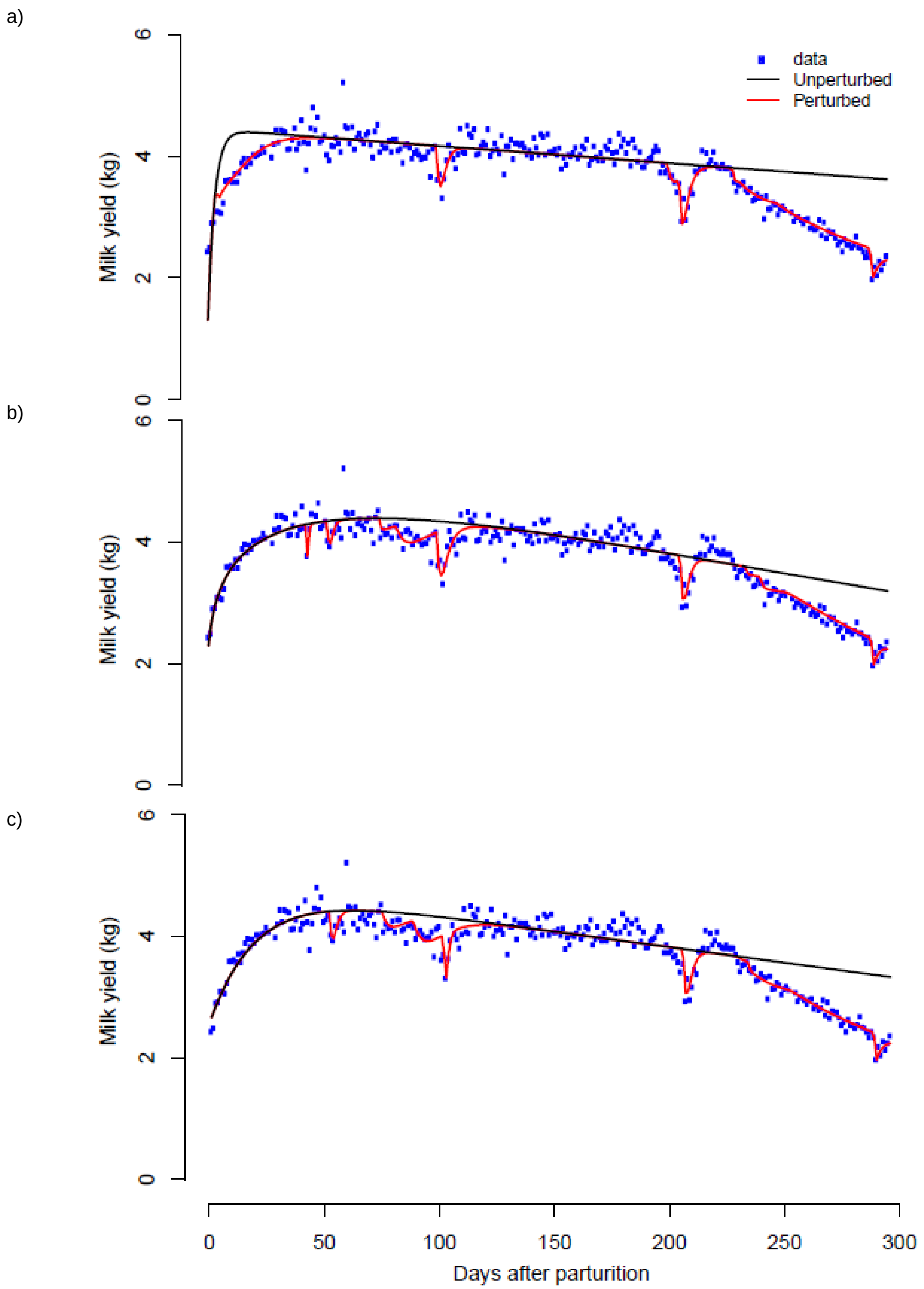


S4. Results with 3 other different lactation models. a : $y_{(t)} = a - b \cdot t - a \cdot e^{-ct}$ (Cobby and Le Du, 1978) ; b : $y_{(t)} = a \cdot b^{ct} \cdot e^{-ct}$ (Dhanoa, 1981) ; c : $y_{(t)} = a + b \cdot e^{-kt} + ct$
